# The Role of Mentoring in Promoting Diversity, Equity, and Inclusion in STEM Education and Research

**DOI:** 10.1101/2021.12.08.471502

**Authors:** Andrea G. Marshall, Zer Vue, Caroline B. Palavicino-Maggio, Elsie C. Spencer, Heather K. Beasley, Edgar Garza-Lopez, Zachary Conley, Kit Neikirk, Sandra A. Murray, Denise Martinez, Jamaine Davis, Lillian Brady, Haysetta D. Shuler, Derrick Morton, Antentor Hinton

## Abstract

Mentoring success is derived from active and respectful listening and the willingness to learn and accept opportunities for personal growth. Mentoring shapes every trainee and their career path in science, technology, engineering, and mathematics (STEM). Productive mentoring relationships cultivate rapport, stimulate moments of introspection, and provide constructive feedback. Effective mentoring in STEM allows trainees, especially underrepresented minorities (URMs), to flourish in welcoming and supportive environments. However, URM trainees often experience inadequate mentoring due to their mentor’s inexperience with URM groups, poor mentorship training, or a lack of understanding of their mentee’s journey. To promote diversity, equity, and inclusion in STEM education and research, it is essential for mentors and mentees to work together with creativity, authenticity, and networking. In this workshop, we will focus on mentees’ perspective on how mentors can enhance their training, professional and career development, and improve their focus. We analyzed data on feedback obtained from students interested in pursuing graduate education who attended a recent workshop. Our results show that despite low initial expectations for the workshop, many students were satisfied with the knowledge they learned. The future of increasing the URM representation in STEM lies in providing adequate community support and mentorship throughout the careers of URM professionals.

## Introduction

Innovation is driven by diversity; as such, it is critical to promote diversity, equity, and inclusion (DEI) within science, technology, engineering, and mathematics (STEM) education to increase innovation (Hofstra *et al*. 2020). Creating a supportive, diverse academic environment is feasible through the leadership of mentors and the support of DEI initiatives. We designed this workshop as an effective tool for improving mentee-mentor relationships with the ultimate goal of increasing retention of underrepresented minorities (URMs) within the academic pipeline (Hinton Jr *et al*. 2020a; Uddin and De Los Reyes 2021).

### The Framework of the Workshop

Effective mentorship is an essential component of every successful STEM career (Hinton Jr *et al*. 2020b; McReynolds *et al*. 2020; Shuler *et al*. 2021; Termini *et al*. 2021b, 2021a). In this workshop, we focused on mentorship and its relationship with DEI. Many trainees, especially URMs, receive subpar mentorship during their education and career. Adequate training in mentorship is essential to drive effective DEI initiatives in STEM. Effective mentors create a supportive, nurturing, and encouraging learning environment while understanding the unique cultural needs and issues of different minority groups (Lewellen-Williams *et al*. 2006). Mentorship is an essential part of the career development of women and minorities in STEM (Kosoko-Lasaki, Sonnino and Voytko 2006; Lewellen-Williams *et al*. 2006). Despite a lack of URM mentors in STEM, well-represented mentors can provide be effective mentorship with adequate training (Lewellen-Williams *et al*. 2006).

#### Fostering Creativity to Promote DEI Initiatives

One simple way for mentors to aid mentees in the development of their DEI portfolio is to incorporate the mentees into their active work. Mentors can help drive DEI at their institution by providing URM trainees additional experience, recognition, and advancement within the STEM pipeline (Hinton Jr *et al*. 2020a; Jayabalan *et al*. 2021). For example, mentors participating in educational outreach at local schools can invite mentees to provide opportunities for hands-on mentoring, leadership, teaching, and communication exposure. Mentors can also be a supportive guide if mentees have ideas for driving DEI initiatives. The expertise that is required to maintain a successful research career is highly transferable into DEI. For example, teamwork, critical thinking, planning, and budgeting are valuable skills. However, the excitement of a proposed idea can overshadow the amount of work involved in creating new DEI opportunities. Mentors can assist in establishing realistic boundaries that enable mentees to have a good work-life balance by setting realistic goals around research, education, and outreach involvement. Thus, mentors have the opportunity to teach mentees the power of saying “no” and delegation (Hinton *et al*. 2020). An effective mentor can help new URM trainees learn essential lifelong skills by teaching them to set realistic goals, how to maintain a work-life balance, and the rewards of involvement and leadership.

#### Identifying a Need: Authenticity

At the core of many DEI initiatives is the desire to drive authentic and meaningful change, which requires a deep understanding of the culture and obstacles faced by URM groups. In some cases, the need may be self-evident. For example, Black scientists are less likely to receive grant funding when compared with their white counterparts with comparable CVs (Ginther *et al*. 2011; Dzirasa 2020; Platt 2020; Stevens *et al*. 2021). Often Black and women scientists face additional challenges when applying for academic jobs (Moss-Racusin *et al*. 2012; Reuben, Sapienza and Zingales 2014; Eaton *et al*. 2020; Hofstra *et al*. 2020). However, other cases may require additional work to identify inequities. In these instances, it is important to approach these communities from a place of authenticity. This requires cultural humility or cultural competency, which involves recognizing the humanity in the population one seeks to serve (Cheng 2007; Foronda 2020). Spending time in getting to know a community’s leaders and trainees demonstrates authenticity and helps build credibility, while also offering first-hand experience in how to address inequities. However, even if a person identified a need and has the resources to assist, they should be able to recognize when to pass a project to a better-suited individual.

Based on these concepts, we tested how students perceived the information and whether they could apply it to their career development and individual development plan (IDP). In this particular questionnaire, we used four questions to gauge interest. The questions consisted of a 10-point scale that was based on rating the following concepts: overall presentation, how to give presentation support team, verbal and nonverbal communication skills, and networking.

#### Content of the Workshop

In this workshop, mentees learned how to fortify their mentor-mentee relationships (Hinton Jr *et al*. 2020b, 2020a; Shuler *et al*. 2021). Cultivating a holistic mentoring relationship based on mutual trust and respect can help foster a stronger, long-term mentor-mentee relationship. This workshop also taught mentees how to become involved in DEI while being successful in science, as well as how to properly communicate DEI interests to a mentor. In addition, this workshop also instructed mentors and mentees effective science communication strategies for gaining public trust and understanding, which includes acting properly to gain credibility over time (Termini *et al*. 2021a). Additionally, this workshop taught participants how to develop DEI initiatives, as well as offered general advice for building long-lasting professional, career, and mentor-mentee relationships. Developing DEI initiatives involves the same skills involved in scientific research: making observations, identifying problems, analyzing outcomes, communicating findings, and presenting solutions. Importantly, this workshop discussed the importance of organizing and planning DEI initiatives, including large-scale efforts. Notably, participants are taught that DEI initiatives are not one-and-done events but require longer-term follow-up to ensure a lasting impact. Long-term involvement displays authenticity and a commitment to building enduring change, which is essential for gaining trust in a community. The workshop also discussed how to replicate STEM education programming initiatives and how that may be used to leverage collaborative mentoring and forge networks. This includes identifying local resources and working with like-minded individuals who may broaden their impact, including community leaders. We also discussed the importance of unity and humility in DEI, by advocating for diverse groups and/or stepping aside for better-suited individuals to manage an initiative when appropriate. Lastly, we covered activities around STEM education. The beginning of the workshop focused heavily on mentorship but then focused on DEI and STEM education efforts. Participants were exposed to the notion of professionalism on social media, regardless of political orientation, as well as cultural competency, which is the ability to work and emphasize with a wide range of individuals and cultures (Cheng 2007; Powell Sears 2012).

### The Necessity of Effective Mentoring in STEM

Traditional STEM mentorship is focused on an individual-centered, information-based, cooperative relationship built on passing on knowledge and experience between a senior to junior researcher. Mentors are entrusted advisors or trainers on various issues including goals, educational and career development, and personal challenges (Hinton Jr *et al*. 2020b; National Academies of Sciences 2020; Shuler *et al*. 2021). Although mentors are often thought of as older and more experienced professionals centered around a career-based apprenticeship, mentors can be of any age. Academic research is fueled by the efforts of trainees at all career stages. Fostering the development of young scientists, especially URMs, is essential for the future of STEM (Hinton Jr *et al*. 2020b; National Academies of Sciences 2020; Shuler *et al*. 2021).

In this workshop, we not only go over the various types of mentoring and the challenges faced by new mentors, we also lay out a blueprint for involving trainees in the development of academic DEI articles. DEI has become a topic of increased focus for many academic institutions in recent years. Authentic relationships and the inclusion of trainees from diverse backgrounds are important for the development of proper DEI-associated training. This workshop also discussed how to use various types of outreach opportunities to provide trainees with the toolkit to excel in STEM; how to gauge the accuracy of scientific information on social media; and how to engage with the general public as a scientist (Ross *et al*. 2012; Streeter 2014; Madakam and Tripathi 2021).

The workshop highlighted the importance of DEI involvement. We challenge mentors and mentees to work together to raise awareness on URM issues. This includes engaging the public by working with different communities within and outside their institution, working with different STEM resources to build rapport, encouraging and developing young scientists, and using effective educational programming to help the advancement of young scholars. Furthermore, we strongly encourage the use of online resources, such as the National Research Mentoring Network, to further enhance the mentoring experience (Sorkness *et al*. 2017; McReynolds *et al*. 2020).

Effective mentorship involves having the willingness to show and lead by example. We suggest that mentors use the supplemental PowerPoint as a template to develop and refine new personalized career development opportunities with their labs and mentees. As such, we encourage mentors to engage in DEI activities in addition to research endeavors. Despite concerns regarding time management, DEI involvement does not harm the chances of obtaining and advancing one’s academic and research career. Publishing academic DEI articles increases the chances of obtaining an academic faculty position (Brandt *et al*. 2021). This suggests that diversity breathes higher levels of innovation that is necessary for success (Hofstra *et al*. 2020).

## Methods

Students from Winston-Salem State University attended a 90-minute virtual workshop and Q&A session. The workshop targeted undergraduates interested in pursuing graduate education to view how the workshop altered their perspective on future academic aspirations. Before and following the workshop, students completed optional, anonymous questionnaires to measure their knowledge about the topics presented in the workshop (Table 1). We analyzed the data using box and whisker plots. Individual answers are represented by circles, standard errors are represented by error bars, and medians are represented by the red centerline. Statistical significance was determined using nonparametric Wilcoxon matched-pairs signed-rank tests. Differences were considered statistically significant when P values were less than 0.05. ****P < 0.0001; ***P < 0.001; **P < 0.01; *P < 0.05; NS, not significant P > 0.05; Data center line, median; boxes, first and third quartiles; whiskers, range; circles represent individual values.

**Table 1.** Pre- and post-workshop evaluations.

| Post-workshop survey questions | Post-workshop survey questions |
| --- | --- |
| On a scale of 1-10, do you think the presentation will keep you well informed? | On a scale of 1-10, how did you like the presentation? |
| On a scale of 1-10, how much do you know about graduate school or professional? | On a scale of 1-10, how much do you know about graduate school or professional school? |
| On a scale of 1-10, how well are you prepared for graduate school before this talk? | On a scale of 1-10, do you feel prepared for graduate school after this talk? |
| On a scale of 1-10, how well do you think the presentation will prepare you for graduate school? | On a scale of 1-10, did the presentation prepare you for graduate school? |
| On a scale of 1-10, how well do you think the workshop will improve your confidence to apply to a professional or graduate school? | On a scale of 1-10, how much do you think the talk helped you improve your confidence to apply to a professional or graduate school? |
| On a scale of 1-10, how well do you think the talk will improve your networking skills? | On a scale of 1-10, how much do you think the talk helped you improve your networking skills? |
| On a scale of 1-10, do you think the talk will improve your understanding of support teams? | On a scale of 1-10, do you think the talk improved your understanding of support teams? |
| On a scale of 1-10, do you think the talk will improve your learning style? | On a scale of 1-10, do you think the talk helped you improve your learning style? |
| On a scale of 1-10, do you think the talk will improve your verbal and non-verbal? | On a scale of 1-10, do you think the talk helped you improve your verbal and non-verbal communication? |

## Results

Pre-workshop questionnaire responses revealed a distinct pattern of students with low expectations. Most students reported not seeing much value in the workshop, as well as a lack of knowledge and confidence about applying to graduate school (Figure 1, pre-workshop). It is unclear if these expectations were influenced by previous experience or simply lack of knowledge. Post-workshop results indicated significant (P < 0.0001) improvement in all categories, including information, preparedness, and confidence in applying to graduate school. Using a 1-10 scale, with 10 being the most favorable response, response scores nearly doubled in all categories. Scores increased by a median of 6 points for the informativeness of the workshop (Figure 1A), suggesting the workshop was more informative than students expected. Similarly, there was a median 5-point increase with regards to students’ knowledge of graduate school (Figure 1B), This is likely linked to the workshop helping to inform students about graduate school, indicated by a large median 7-point increase (Figure 1C). Finally, participating students felt more prepared and confident in applying for graduate school, suggested by a median 5- and 6-point increase, respectively (Figure 1D and E). These results indicated that the students were initially hesitant about the effectiveness of the workshop but found it worthwhile.

**Figure 1.**
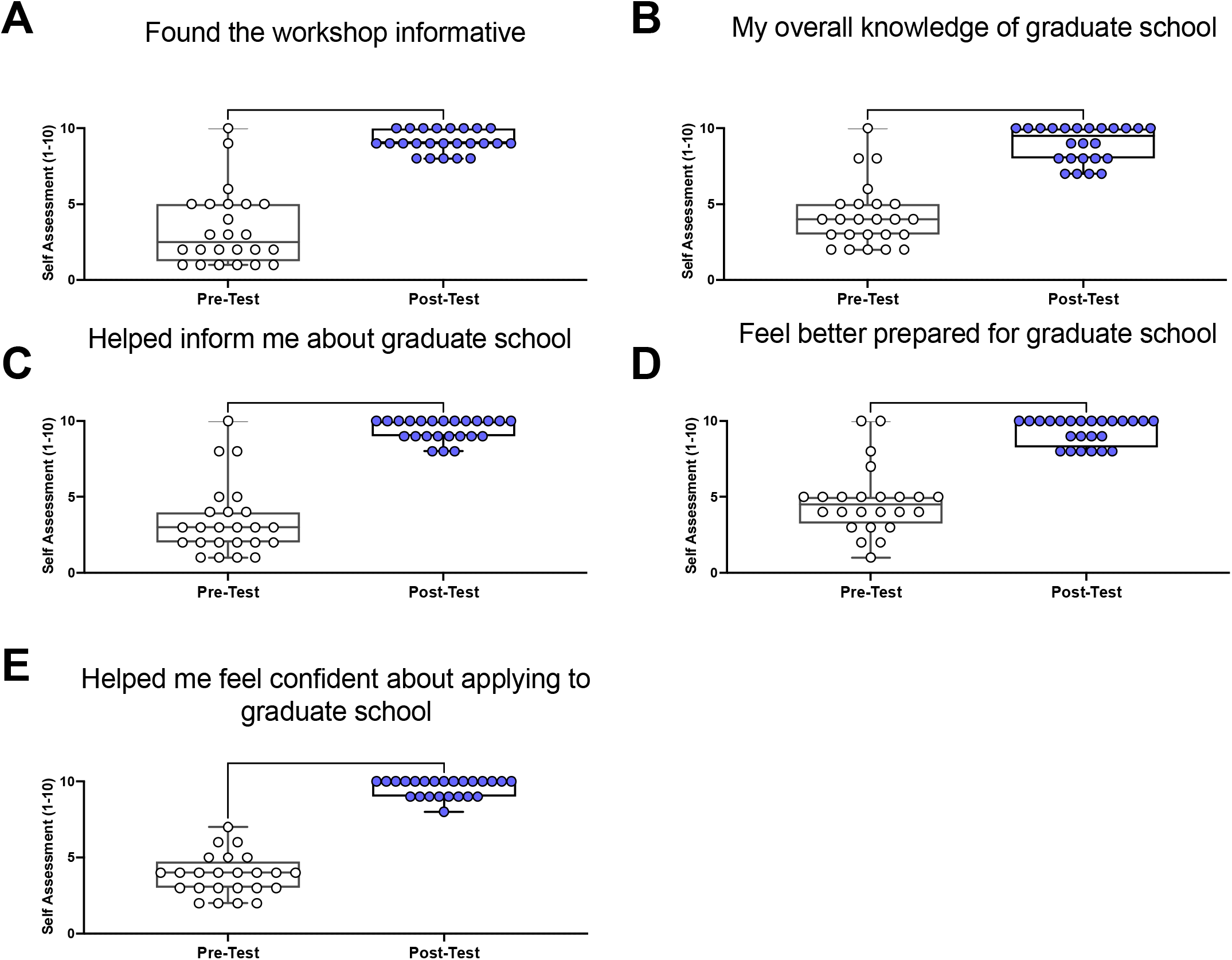
Results from pre- and post-workshop evaluations for questions on their overall knowledge of graduate school.

We found similar results regarding questions related to the mentee aspect of mentorship and the applicability of the content. Pre-workshop questions related to networking skills, support teams, learning style, and communication skills found students feeling ill-informed (Figure 1 & 2, pre-workshop). We observed a strong significant increase in scores for all mentoring questions following the workshop, indicating students felt more informed about mentoring relationships. Although students did not expect this workshop to aid in networking skills (average initial score: 3.4), the average score increased by 5.8 (Figure 2A). Similarly, the students’ knowledge of support teams rose by 5.3 following the workshop (Figure 2B). This workshop focused on learning styles. Students were moderate to moderately knowledgeable of learning styles before the workshop (average: 4.8); however, this average increased to 9.9 post-workshop (Figure 2C). Students also reported that the workshop helped improve their verbal and non-verbal communication skills; averages rose from 4.6 to 9.3 (Figure 2D). Altogether, student expectations of the workshop ranged from low to moderate; however, they found the workshop informative, suggested by increased post-workshop scores. These findings suggested that the workshop was effective in encouraging students to consider graduate school, as well as teaching them the skills required in having a strong mentor-mentee relationship.

**Figure 2.**
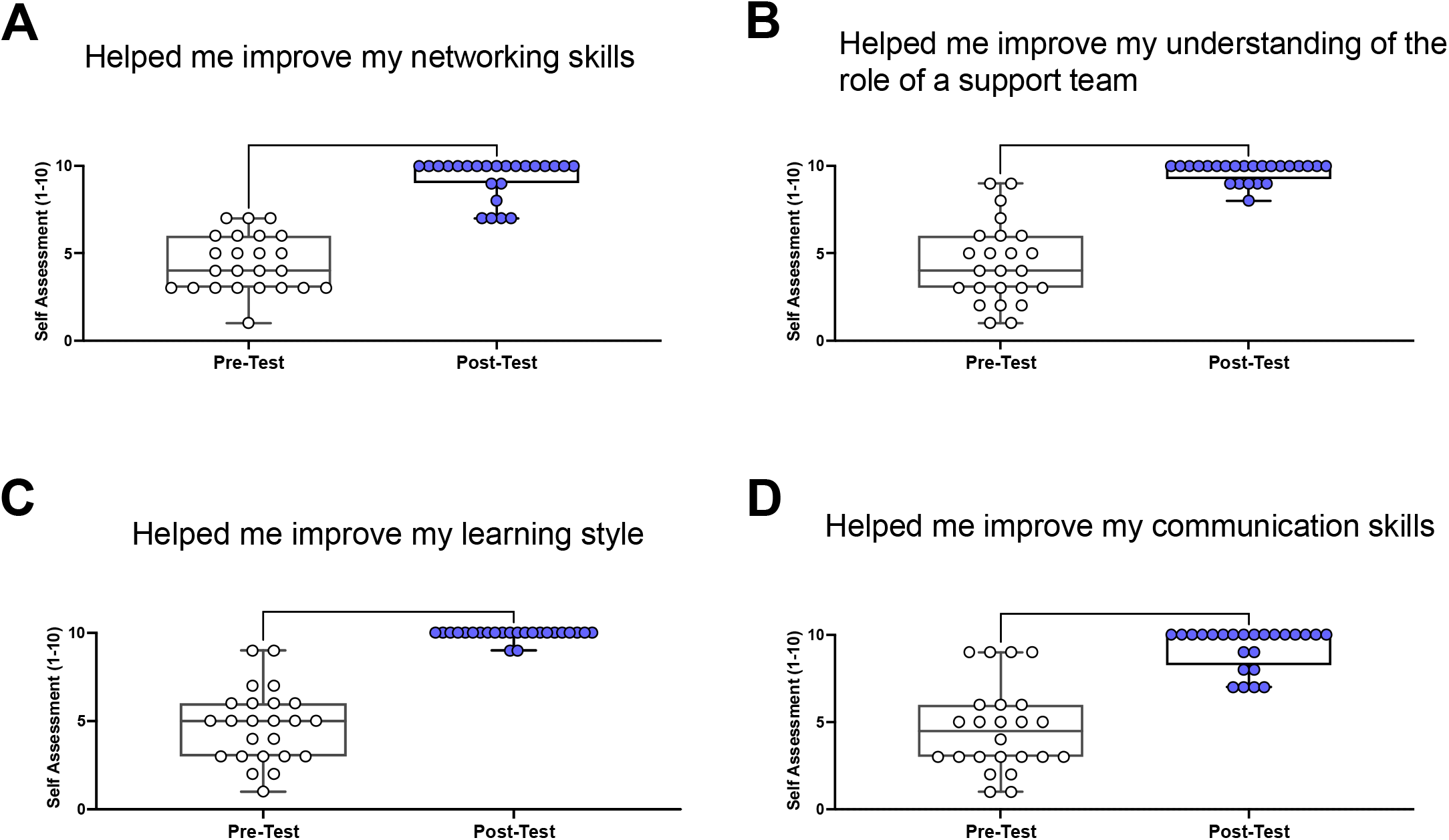
Results from pre- and post-workshop evaluations on how well the workshop helped inform them of skills needed to navigate successfully in graduate school.

## Discussion

Our study demonstrates the need to educate academic professionals, faculty, and students about mentorship and support teams while encouraging them to work together.

Our data supports the need to learn networking skills. Mentors can share their networking expertise and help trainees build a network. Despite being essential to career development, networking skills are tough to develop as many trainees receive few opportunities, such as conferences and meetings, to practice these skills (Streeter 2014). These networking opportunities allow trainees to expand their mentorship away from official settings, finding them within their family, organizations, religious settings, and even among peers. Thus, students can apply the knowledge gained by this workshop to develop mentorships at all stages of their careers.

Although academic institutions are now understanding the mutual relationship between diversity and innovation, it is also important to cultivate inclusive environments (Hofstra *et al*. 2020). Our findings suggest that students are supportive of having opportunities involving diversity and inclusion to enrich their training experience (Fig. 1, 2). Workshops can be used as a necessary resource to create environments that support DEI, diverse thinking, and effective mentoring.

Although there is a developing interest in encouraging students to become involved in DEI, there are misunderstandings about the time commitment needed to do a good job. We suggest supporting and listening to students interested in STEM and DEI outreach to get the proper resources to establish a network of DEI professionals. Our findings also suggest the need for consistent conversations that span from mentee-mentor relationships to departments, dean offices, and the institution to encourage continued efforts. This will allow trainees to get the training they need even when an effective mentor is not available.

## Availability of Data and Materials

PowerPoint presentations of the workshop are available in English and Spanish. Survey data is available upon reasonable request.

## Acknowledgements

We thank the 24 students who participated in our survey.

## Data and materials availability

All data are available in the main text or the supplementary materials.

## Ethics Declaration, Project Title

Promoting Engagement in science for underrepresented Ethnic and Racial minorities (P.E.E.R), 21-MortonD-HSR-SOM-01, Kaiser Foundation Research Institute FWA: FWA00002344

## Ethics Approval and consent to participate

Yes

## Consent for publication

Yes

